# The emergence of inter-clade hybrid SARS-CoV-2 lineages revealed by 2D nucleotide variation mapping

**DOI:** 10.1101/2020.10.13.338038

**Authors:** Hai-Long Wang

## Abstract

I performed whole-genome sequencing on SARS-CoV-2 collected from COVID-19 samples at Mayo Clinic Rochester in mid-April, 2020, generated 85 consensus genome sequences and compared them to other genome sequences collected worldwide. I proposed a novel illustrating method using a 2D map to display populations of co-occurring nucleotide variants for intra- and inter-viral clades. This method is highly advantageous for the new era of “big-data” when high-throughput sequencing is becoming readily available. Using this method, I revealed the emergence of inter-clade hybrid SARS-CoV-2 lineages that are potentially caused by homologous genetic recombination.

## INTRODUCTION

The COVID-19 pandemic caused unprecedented disruption to populations worldwide. The causative agent is the SARS-CoV-2 virus, a single-stranded positive-sense RNA virus from the beta-coronavirus genus with ~30 kb in genome length^1,2^. Although general testing for this novel virus is rapidly expanding, targeted sequencing approaches have attracted more attention for the following reasons: a) The SARs-CoV-2 virus has evolved into mutant strains with different clinical complications. Monitoring the spread of specific strains during an outbreak can help inform intervention methods and their impacts. b) Vaccine efficacy can be monitored by tracking the prevalence and distribution of key genes involved in vaccine development and deployment, and by determining the viral mutation rates to inform a vaccine’s longevity and its interactions with the human immune system ^3–5^. Today, there are over 100,000 genome deposits on GISAID (the Global Initiative on Sharing All Influenza Data) and over 30,000 on NCBI (National Center for Biotechnology Information), reflecting a worldwide effort to fight against the virulent pathogen. The wealth of SARS-CoV-2 genome sequences provides a rare opportunity for both clinical and basic scientists to investigate its evolution mechanism. On the other hand, outsized collections of genome sequences pose a computational challenge to bioinformatics analysis. Currently, the most adopted method for illustrating species evolution, the phylogenetic analysis, is inadequate for discerning fast-evolving mutations and recombination. New methods for mapping mutations within the same genome are needed for the impending “big-data” era. I propose a two-dimensional nucleotide variant mapping that illustrates populations of co-occurring mutations within the same genome. This plotting method is not limited by the increasing number of examined sequences. Thus, it allows a direct comparison between genome collections from different geographic regions or the same region but at a different time. Furthermore, it reveals the emergence of inter-clade hybrid lineages.

## RESULTS

### 1. SARS-CoV-2 genome sequencing

As part of Mayo Clinic’s efforts to fight against COVID-19, I performed viral genome sequencing using the third-generation sequencing techniques developed by Oxford Nanopore Technology (ONT). I received 103 positive COVID-19 samples of RNA extracts collected by the Department of Laboratory Medicine & Pathology at Mayo Clinic Rochester from April 20, 2020 through April 29, 2020. Following the ARTIC network’s amplicon sequencing protocol^6^ (see method for details), I succeeded in collecting 85 genome sequences.

A phylogenetic tree, presented in Figure 1, compares these genome sequences to a reference genome (NC_045512.2). According to the latest SARS-CoV-2 classification^7^, there are six major viral clades (named L, S,V, G, GH and GR, respectively) represented by their unique sets of single nucleotide polymorphisms (SNPs). Clade “L” is the reference genome (NC_045512.2) found in China; clade “S” is named after the mutation in ORF8:L84S (with a transition mutation of T28144C and co-occurring with another transition mutation of C8782T); clade “V” is named after the mutation in ORF3a:G251V (with a transversion mutation of G26144T and co-occurring with another transversion mutation of G11083T); clade “G” is named after the D614G mutation in the spike protein (with a transition mutation of A23403G and co-occurring with other three transition mutations of C241T, C3037T, C14408T). Two derivative clades from the G clade are the “GH” clade characterized by a mutation ORF3a:Q57H (G25563T), and the “GR” clade characterized by the trinucleotide mutations in the nucleocapsid gene, inducing G28881A, G28882A and G28883C. From these 85 genome sequences, 44 belong to the S clade (in magenta), 1 to the V clade (in red), 18 to the G clade (in green), 18 again to the GH clade (in cyan), and 4 to the GR clade (in orange). The same coloring scheme for distinct clades will be used throughout the paper for convenience.

**Figure 1.**
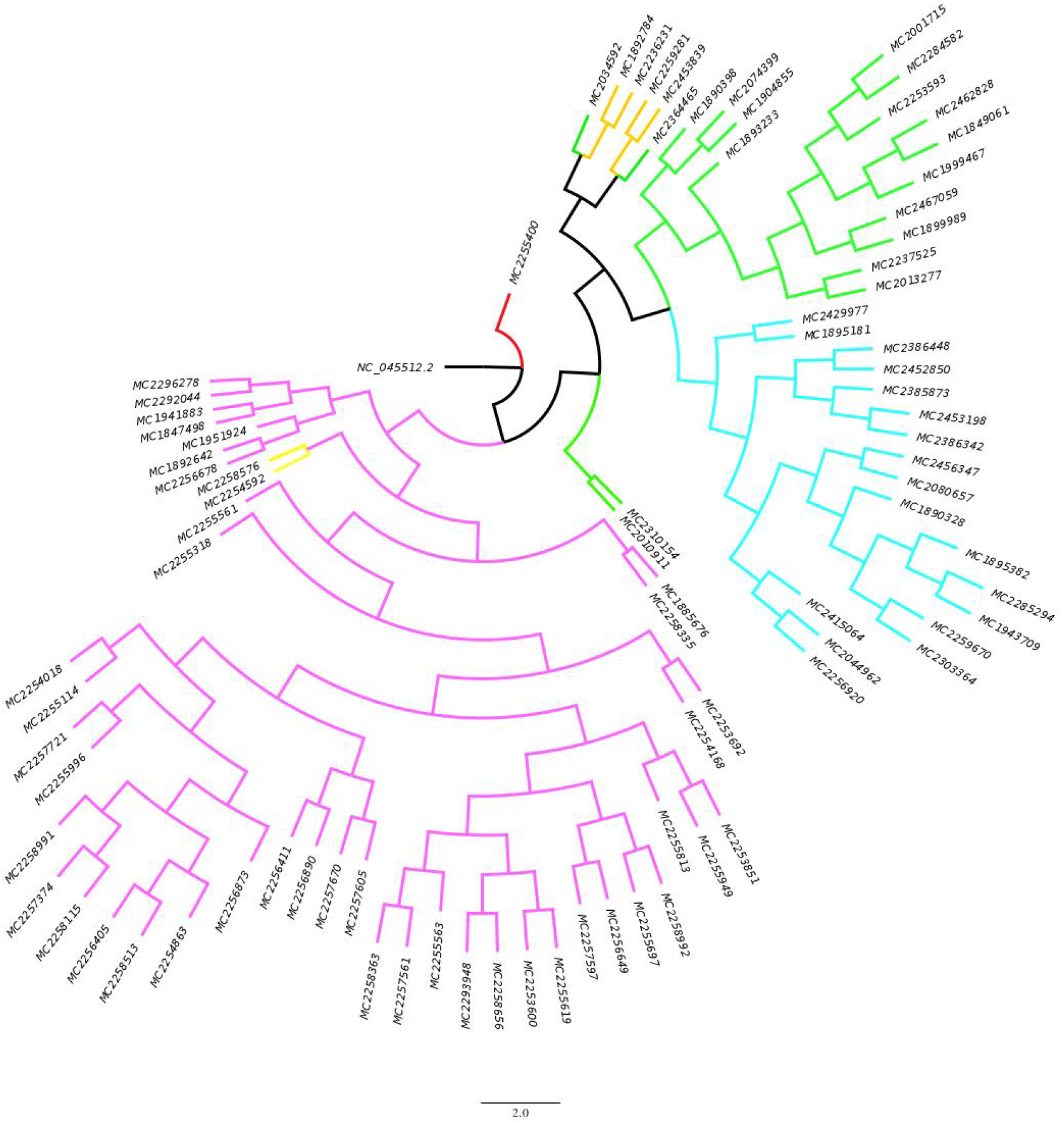
A phylogenetic tree compares 85 genome sequences to the reference genome (NC_045512.2). Distinct clades are indicated with colored lines: 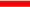 V clade, 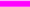 S clade, 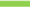 G clade, 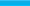 GH clade, 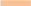 GR clade. The picture was generated with the FigTree software.

Profiling SNPs is another way to show frequencies of mutations within a clade. Shown in Figure 2 are profiles of two dominant clades, S (Fig 2A) and G (Fig 2B). High-profile SNPs are those with top heights representing high frequency at special genome loci. Counts of less than two were omitted to reduce the background noise caused by low-profile *de novo* mutations. Being unique to distinct clades, high-profile SNPs are considered as clade-defining SNPs.

**Figure 2.**
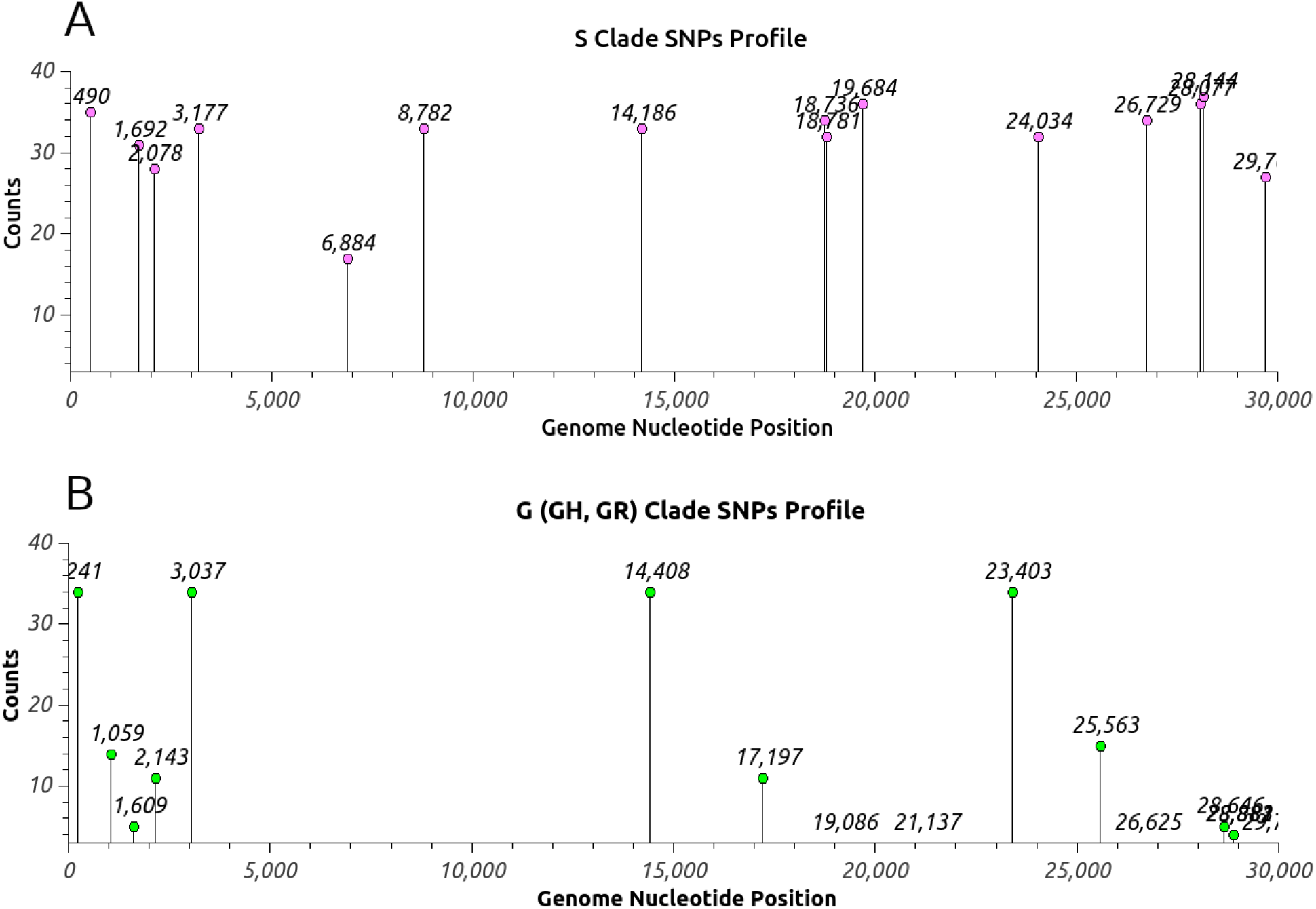
SNP profiles for the S clade (A) and the G clade (B) from 85 genome sequences. The height of each dot represents a counted number of a particular variant at the nucleotide position from all genome sequences. Counts of less than two are omitted to reduce the background noise caused by low-profile SNPs.

### 2. 2D nucleotide variation analysis

The phylogenetic analysis, as commonly used to display evolutionary relationships among species, employs algorithms to calculate the evolutionary difference between two species and uses estimated distances to draw branching patterns that display how the species evolved from a common ancestor. Although a phylogenetic tree is appealing, it traditionally ignores the recombination event that is another major driving force in evolution and causes closely related species to have different phylogenetic histories.

SARS-CoV-2 is an RNA virus with limited genome length, but its high mutation rate and homologous genetic recombination nonetheless gave rise to exponentially increased variants. Classical studies based on consensus sequences become insufficient to interpret its adaptive potential. Besides, drawing a phylogenetic tree for many species becomes impractical due to constraints in the drawing space (e.g. it was difficult to layout merely 85 sequences in Figure 1). An alternative method is needed for the impending era of “big-data”, when the full wealth of sequencing collections is becoming available. This new method should also demonstrate the links between nucleotide variations on the same genome.

Here, I propose a 2D mapping for co-occurring SNP pairs. In brief, a matrix (n x n) is created in which “n” is the genome length. From each genome sequence, any detected co-occurring SNP pair will be used to increase the value of M_ij_ by 1, where “i” and “j” are the two positions (loci) of nucleotides from that SNP pair. The final matrix is then normalized by dividing each element by the total number of genome sequences. Directly plotting of this matrix into a 2D scaled map will generate a diagonally symmetric pattern that displays the distribution of population percentages for every co-occurring SNP pairs in the whole genome set. However, including all ~30,000 nucleotides in a full-scale image is overwhelming for normal human eyes to process. A simpler format including only a selected set of loci would be a better way to demonstrate the concept. I plotted a bubble map with those 12 clade-defining SNPs (including 241, 3037, 8782, 11083, 14408, 23403, 25563, 26144, 28144, 28881, 28882, and 28883 in the order of their nucleotide position in the genome and colored according to their representing clades). The radius of each bubble is proportional to the corresponding population percentage (see Figure 3).

**Figure 3.**
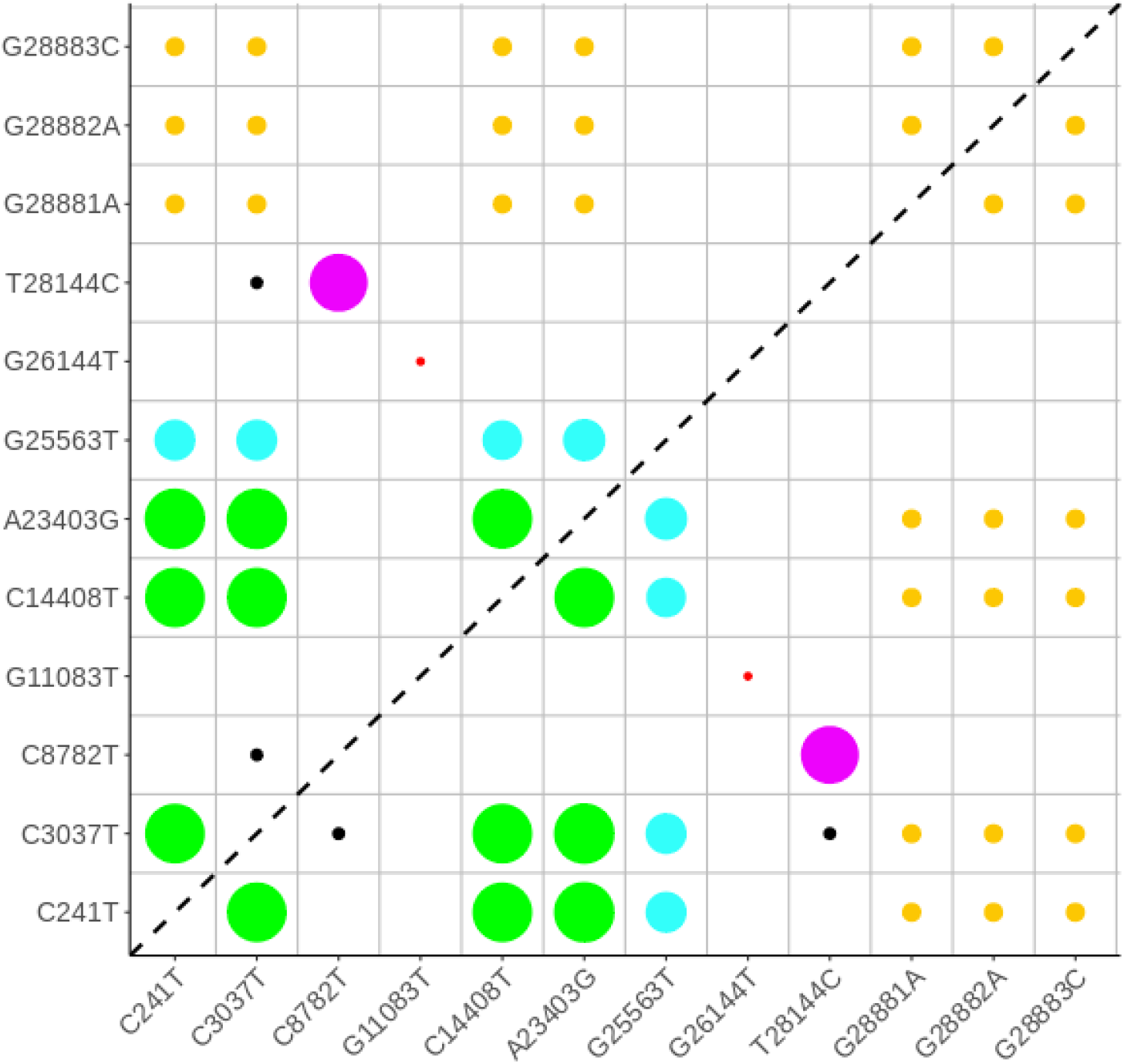
The 2D bobble map of co-occurring SNP pairs from 85 genome sequences. Distinct clades are indicated using a coloring scheme: 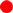 V clade, 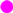 S clade, 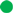 G clade, 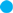 GH clade, 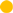 GR clade, 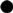 inter-clade. Radii of bubbles are proportional to population percentages of distinct clades. The dashed line reflects the diagonal symmetry of the map. The map was generated by an R script.

In this kind of 2D map, a colored bubble at an intra-clade intersection represents a population of the representing clade; while regions at inter-clade intersections are normally empty, indicating no crossover between distinct clades. In Figure 3, however, there are two (or four, if accounting for the diagonal symmetry) regions belonging to two pairs of nucleotides at (3037, 8782) and (3037, 28144) that are not empty but have small black dots. The locus of 3037 belongs to the G clade and both loci of 8782 and 28144 belong to the S clade. These inter-clade co-occurring SNP pairs are found in two lineages (MC2254592 and MC2258576) that were previously highlighted in the phylogenetic tree of Figure 1.

There are two possibilities for inter-clade crossover events: 1) a *de novo* mutation; 2) homologous genetic recombination. These two lineages both have a total of eight SNPs (for MC2254592: 2078, 3037, 8782, 19684, 24043, 28077, 28144, 28849; for MC2258576: 3037, 3177, 8782, 11897, 19684, 24034, 25505, 28144, colored according to their representative clades). According to the SNP profile of the S clade shown in Figure 2A, the lineage MC2254592 has six high-profile SNPs characteristic to the S clade, while MC2258576 has five, which might be the reason why the phylogenetic tree includes them both in the S clade. Other than the high-profile SNP at 3037 representing the G clade, there is only one low-profile mutation in MC2254592 and two in MC2258576. The small number of low-profile SNPs found in both lineages suggests that the possibility of a *de novo* mutation causing the high-profile SNP at 3037 to occur in an S-clade lineage is very low.

Homologous genetic recombination, on the other hand, is a major evolutionary driving forces found in all kingdoms of living organisms. Viral homologous recombination, discovered in several other RNA viruses, has the potential to create a novel strain with enhanced virulence^8–11^. For the SARS-CoV-2 virus, several accounts of recombination events have been reported^12–14^.

There are several tools, including Simplot^15^, RDP4 ^16^, and PhiPack ^17^, that are specifically designed for detecting homologous recombination. But none of these tools can detect any possible recombination events among 85 genome sequences, possibly due to low-frequency events with few SNPs. For example, the PhiPack was designed to work for low-divergence genomes, it still requires 1~5% of variants to perform statistical analysis. Because of the relatively short period into COVID-19 pandemic, most viral lineages have less than 20 SNPs, which is at only 0.067% of variant percentage. This is why all previously reported recombination events of SARS-Cov-2 have relied on clade-defining SNPs.

The 2D co-occurring variant mapping is a simple way to display inter-clade hybrid lineages, and it can be used to directly compare distributions of populations for intra- and inter-clade from different geographic locations or the same location but at a different time point. Therefore, it is worth to examine genome sequences from around the world that have been deposited into public domains.

### 3. The big picture of homologous recombination

#### 3.1. U.S. domestic regions

I downloaded ~18,000 SARS-CoV-2 genome sequences from NCBI (on September 2^nd^) and used the same criterion as before to search for inter-clade co-occurring SNP pairs (see method for details). I first looked at domestic regions in the US and focused on four states, including New York, Minnesota, California, and Washington, that represent different geographic regions (see Figure 4).

**Figure 4.**
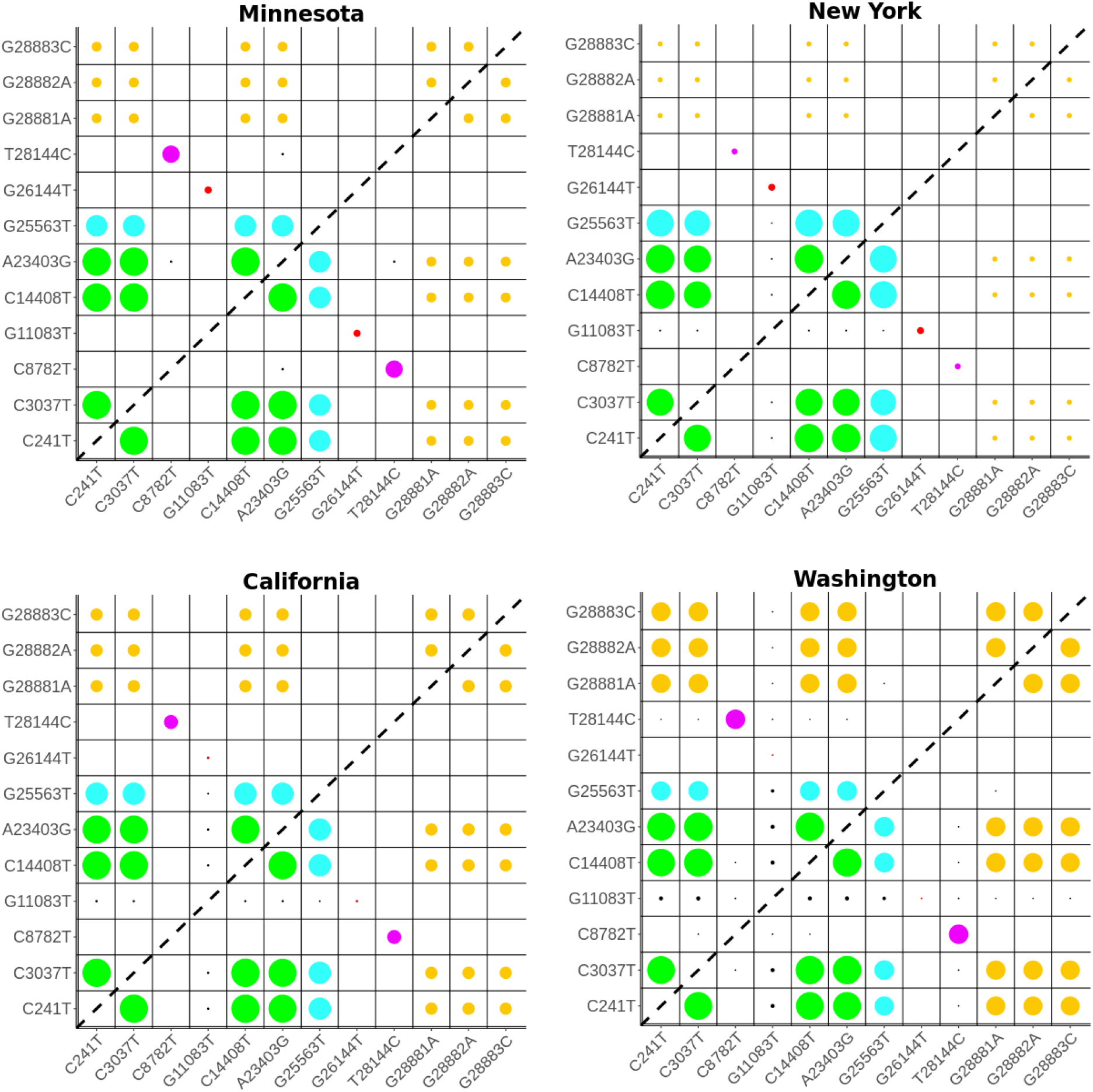
The 2D bobble maps of co-occurring SNP pairs from four states in the US, including Minnesota, New York, California, and Washington. Distinct clades are indicated with color: 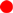 V clade, 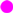 S clade, 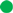 G clade, 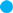 GH clade, 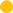 GR clade, 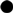 inter-clade. Radii of bubbles are proportional to population percentages of distinct clades. The dashed line reflects the diagonal symmetry of the maps.

The Minnesota genome collection had 124 sequences at the time of this study. Its 2D plot shows a pattern similar to that in Figure 3 (no doubt given that Mayo Clinic Rochester is located in Minnesota), which is dominated by the S and the G clade, followed by a smaller population of the GH clade and an even less-populous GR clade, with the V clade as the least populous. There are also two inter-clade co-occurring SNP pairs at (8782, 23403) and (28144, 23403) between the S and the G clade. Both co-occurring SNP pairs were identified in the same lineage (NCBI access ID: MT920018).

The New York genome collection had 285 sequences, and its 2D plot shows two equally dominant G and GH clades, followed by V, S, and GR clades. Inter-clade co-occurring SNP pairs are found between the locus at 11083 of the V clade and all other representative loci of the G and GH clades. These co-occurring SNP pairs were identified in two lineages (NCBI access ID: MT370869, MT370986).

The California genome collection had 916 sequences, and its 2D plot shows increased differences when compared with that of New York. In California, the largest cluster still belongs to the G clade. Yet, both the S and GR clades are significantly bigger than those found in New York. The V clade is the least-populous. Again, inter-clade co-occurring SNP pairs are found between the 11083 of the V clade and all other representative loci of the G clade and GH clade that correspond to six lineages (NCBI access ID: MT628091, MT628092, MT628093, MT750350, MT750445, and MT806758).

The Washington genome collection had 3,572 sequences, and its 2D plot shows significantly increased populations in the S and the GR clades. Three groups of inter-clade co-occurring SNP pairs are revealed. The first one is between the locus at 11083 from the V clade and all other major clades; the second one is found between the S and the G clades, involving both representative loci (8782 and 28144) of the S clade; the third one is at (25563, 28881/2/3) between the GH and the GR clades. The long list of corresponding lineages can be found in supplemental Table s1. Here, I paid my attention to the second group, in which two representative loci simultaneously crossing over into a distinct clade, which would be stronger evidence for homologous genetic recombination.

Three candidate lineages were identified (also labeled with an underline in Table S1. NCBI access ID: MT834063, MT821583, MT834220). The lineage MT834063 has 11 loci (316, 1526, 5622, 8782, 14408, 17561, 17747, 19839, 23043, 28144 and 29784), whereas the other lineage MT834220 has 9 loci (241, 3037, 8782, 17747, 17858, 28144, 28881, 28882 and 28883), all colored according to distinct clades. In the case of MT834063, it is expected that the genome region between 14408 and 23403 of the G-clade strain was swapped with the same region in an S-clade strain. In the case of MT834220, however, it is expected that the region between 8782 and 28144 of the S-clade strain was swapped with the same region in a GR-clade strain. Also, the absence of the locus at 23043 in MT834220, a signature SNP of all G-(GH and GR) clade lineages, is another strong piece of evidence for homologous genetic recombination. The third lineage of MT821583 is similar to MT834220 but has additional low-profile SNPs.

#### 3.2 International regions

So far, I have demonstrated the usage of 2D co-occurring SNP mapping for detecting inter-clade hybrid lineages of the SARS-CoV-2 virus. Next, I also examined genome collections from international regions and focused on the following four countries: China, Japan, Italy, and Spain, where early COVID-19 outbreaks were detected (see Figure 5).

**Figure 5.**
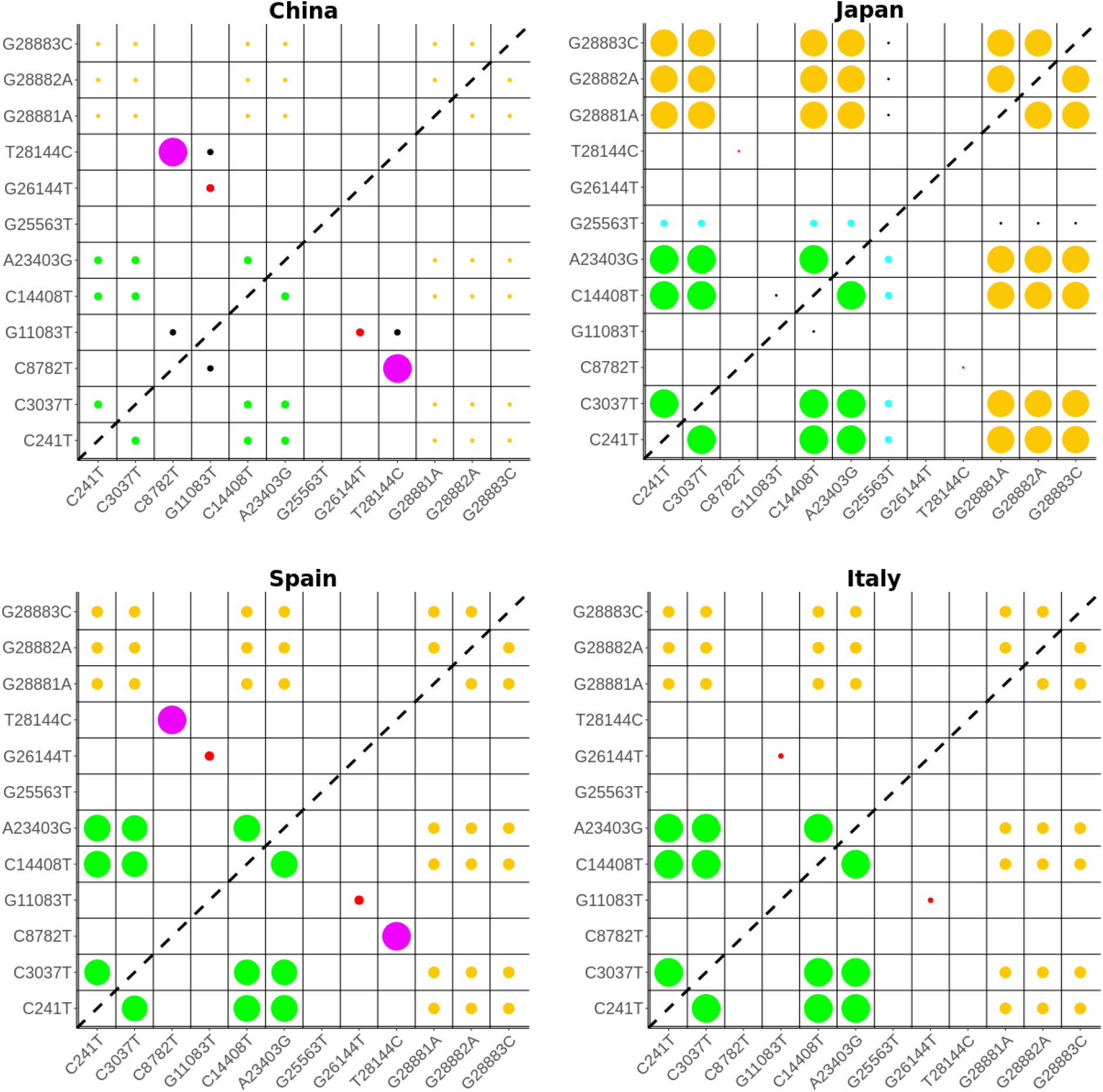
The 2D bobble maps of co-occurring SNP pairs from four countries, including China, Japan, Italy, and Spain. Distinct clades are indicated with color: 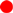 V clade, 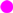 S clade, 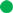 G clade, 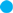 GH clade, 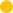 GR clade, 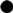 inter-clade. Radii of bubbles are proportional to population percentages of distinct clades. The dashed line reflects the diagonal symmetry of the maps.

2D plots of these four regions are very different. The genome collection from China had 79 sequences with the S clade being the most dominant clade. It is followed by the V, G, and GR clades based on the population size. No GH clade was detected. Two inter-clade co-occurring SNP pairs were found at (8287, 11083) and (28144, 11083), both are between the S and the V clades, corresponding to two lineages (NCBI access ID: MT049951 and MT226610). The genome collection from Japan had 84 sequences with the GR clade as the largest cluster, followed by a smaller GH clade. The S-clade is the least-populous. No viral lineage of the V clade was detected. Two inter-clade co-occurring SNP pairs were found, one at (11083, 14408) between the V and the G clades (NCBI access ID: LC542809), and the other one at (25563, 28881/2/3) between the GH and the GR clades (NCBI access ID: LC556322).

The genome collection from Spain had 37 sequences. Two dominant clusters are the S and G clades, followed by the GR and the V clades. No GH clade was detected. The genome collection from Italy had 24 sequences with the G clade as the largest cluster, followed by the GR and V clades. No GH or S clade was detected. No inter-clade co-occurring SNP pairs were detected in genome collections from these two countries deposited to NCBI on September 2^nd^.

## DISCUSSIONS

The COVID-19 pandemic caused unprecedented disruption to populations worldwide. Fighting the causative agent is the indisputable duty of everyone who has the relevant knowledge and skills. Sequencing the viral genome can help characterize the evolutional changes of the virus and help public health authorities understand the identity, whether it is changing, and how it is being transmitted. I procured a MinION device in January originally for another sequencing project. My experiences learned during this project are worth further discussions. I will mainly focus on the following four topics: the use of nanopore sequencing technology, pros and cons of the phylogenetic analysis, homologous genetic recombination, and the quasispecies model of RNA viruses.

### 1. Nanopore sequencing of the SARS-CoV-2 virus

Oxford Nanopore Technology gained attention for its long-reading sequencing technology, low cost, and portable capability. Many researchers are using the MinION device (costing $1,000) as basic laboratory equipment for high-throughput sequencing needs. The system has been previously used in other outbreak situations, such as EBOV ^18^ and ZIKV ^19^. Early in March, the ARTIC network opened a specific sequencing protocol for COVID-19, including a set of primers, laboratory materials, and related tools for bioinformatics analyses. The protocol relies on direct amplification of the virus using tiled and multiplexed primers, and it has been widely adopted by many groups around the world^20,21^.

The cost of two flowcells (>$500/each) and other reagents is about $2,000. A rough estimation of the material cost per sample is about $20~$30. The bottleneck for the study is preparing the library before sequencing. It took me a whole week to prepare 103 samples, which were all processed manually. I made some mistakes due to a lack of experience, so only 85 consensus sequences were finally achieved. The library preparation process can be easily scaled up with a liquid handling robot. With 1~2 hours of data acquisition, the MinION device usually generates one million reads, enough to provide results for collecting final consensus sequences. The feature of live-basecalling is necessary when using the RAMPART tool ^22^ to monitor the on-going progress during sampling and to check whether all amplicons have been sampled. The final stage for bioinformatics analysis was handled by the nextflow COVID-19 workflow that includes read-quality score filtering, reference alignment, and consensus sequence generation, which only takes about twenty minutes. This low cost, decentralized sequencing can be set up quickly for flexible, portable, and scalable applications.

### 2. Limitations of phylogenetic analysis

A phylogenetic tree is a diagram used to illustrate phylogeny, the history of evolution, in reference to lines of descent and relationships among broad groups of organisms. The branching patterns in a phylogenetic tree reflect how species or other groups evolved from a common ancestor. The methodology has been used increasingly in diverse areas of biological science, especially for tracing viral evolution at the early stages of the outbreak in an epidemic like COVID-19 ^3,21,23,24^.

However, there are inherent limitations to a phylogenetic tree. First, traditional methods of phylogeny estimation, such as maximum parsimony, minimum evolution, or maximum likelihood, all assume that a single evolutionary history underlies the sequences. Second, the viral network presented in the tree diagram is merely a snapshot of the early stage of the viral spreading before the phylogeny becomes obscured by subsequent migration and mutation. When the number of species in the group increases, drawing a tree diagram becomes impractical. Third, commonly used methods in phylogeny, such as the pairwise distance method, do not capture the effects of homologous recombination that imply how different parts of the sequence could have separate phylogenetic histories and are not related by a single phylogenetic tree^25^. Though the ability to detect recombination is limited, ignoring recombination in tree-based analysis could lead to artifacts^26,27^.

### 3. Homologous recombination

In the nature of evolution, viruses are continuously changing as a result of genetic selection through two principal mechanisms: 1) *de novo* mutation that occurs when an error is incorporated in the genome and causes subtle genetic changes; 2) recombination that happens when co-infecting viruses swap genetic information and cause major genetic changes^28^.

There are two mechanisms of viral recombination: 1) the independent assortment, where viruses with multipartite genomes trade unlinked, and assorted random segments during replication; 2) the incompletely linked gene, where viruses trade linked genes residing on the same piece of nucleic acid. Either mechanism can produce new viral serotypes with altered virulence^8^.

Recombination has been shown to occur in several positive-sense single-stranded RNA virus groups, such as retroviruses, picornaviruses, and coronaviruses. It is currently believed that recombination in coronaviruses occurs by a copy-choice mechanism^28^, in which the viral RNA polymerase binds to only a few bases of the template RNA. The weak interaction permits the polymerase to disassociate from the original template and then associate with a new template RNA strand. The efficiency of this mechanism of recombination is relatively low.

Viral recombination generates novel progeny viruses that express new antigenic characteristics, with new surface proteins to infect previously resistant individuals, or with novel combinations of proteins to infect new cells in the original host. On the other side, viral recombination has also been used to create new vaccines. Vaccinia virus strains carrying genes coding for viral antigens but low virulence has been produced to stimulate specific antibody production by the host, resulting in the protection of the host from the immunogen. Therefore, characterizing viral recombination will not only help health management teams effectively contain emerging viruses with enhanced virulence, but also support virologists in creating new vaccines.

### 4. Quasispecies model and 2D co-occurring variants analysis

Due to their limited genome length and higher mutation rate, many RNA viruses have genetically diverse populations known as quasispecies^29^, meaning a highly diverse replicating population reaching equilibrium in the level of diversity.

In traditional population genetics, a given variant in a population is approximated by its fitness. However, the quasispecies model suggests that variants are “coupled” in sequence space. A low fitness variant can thereby be maintained at a higher than expected frequency because it is coupled to a well-represented, higher fitness genotype in sequence space^30^.

Instead of focusing on ancestor-descendant relationships, the quasispecies model puts more weight on co-occurring variants within the same genome and their corresponding populations, which is exactly the purpose of this 2D mapping for co-occurring variants. The emergence of inter-clade hybrid SARS-CoV-2 lineages indicated that the causative agent of COVID-19 has entered a phase of quasispecies.

While tracing the path of viral infection is still important, estimating the level of viral diversity is also urgently needed. In order to survive, a virus must be diverse enough to rapidly adapt to changing environments without losing fitness during the passage from host to host. The level of diversity varies depending on the host environment and is specific to the virus-host combination. The differences in viral clade populations from separate regions are clearly displayed in Figures 4 and 5.

Predictions from the quasispecies theory have profound implications for the understanding of the viral disease. The 2D plotting method for co-occurring variants is a useful tool for a longitudinal study to monitor the level of diversity from a region with certain environments, include the ethnogenesis population, the policies enforced by local health authorities, and viral immunity in the area.

In summary, I performed whole-genome sequencing of the SARS-CoV-2 virus and compared those genome sequences with other samples collected worldwide. I proposed a 2D plotting method to illustrate co-occurring variants, as it has several advantages: 1) it directly shows the population of coupled mutations without any statistical manipulation. 2) it is suitable to display big data with no limit on the size of the genome collection (actually the more the better); 3) it is flexible in regards to the plotting technique. Even though the full picture with the whole range of genome length contains an overwhelming amount of information, a simplified snapshot could be generated from a subset of the sequence space; 4) it helps identify the emergence of inter-clade hybrid viral strains that have both positive and negative potential impacts on all of us.

## METHODS

### Genome sequencing

Specimen collection and testing of individuals with suspected COVID-19 at Mayo Clinic Rochester were screened for SARS-CoV-2 infections as per the CDC protocol. 103 samples of COVID-19 positive RNA extracts collected between April 20, 2020 and April 29, 2020 were provided by the Department of Laboratory Medicine & Pathology under Mayo Clinic’s approved protocol for COVID-19 study.

Detailed whole genome sequencing and analysis methodologies came from the ARTIC network^6^. Briefly, cDNA synthesis was performed with a SuperScript IV First Strand Synthesis Kit (Thermo) according to manufacturer specifications. Direct amplification of the viral genome cDNA was performed using multiplexed primer pools per protocols provided by the ARTIC Network (V3). The sequencing library preparation protocol was adapted from the ARTIC Network’s protocol for ONT “PCR tiling of COVID-19 virus”. Adapter ligation and cleanup were performed using the ONT’s Ligation Sequencing kit (SQK-LSK109) according to manufacturer specifications. Libraries were sequenced on ONT’s MinION device using Type R9.4.1 flow cells (FLO-MIN106D). Raw sequencing data were acquired using ONT’s MinKnow software (3.6.5) running on a custom-built Dell Precision T5600 desktop computer equipped with an NVIDIA RTX 2080Ti graphic processing unit. Fast live-basecalling was enabled for barcode detections.

High-accuracy basecalling of nanopore reads was later performed using Guppy v3.4.5 on an HP Prodesk830 computer equipped with an NVIDIA RTX 2060 GPU. Automatic genome analyses were performed with the ARTIC COVID-19 nextflow software package for read-quality score filtering, reference alignment, and consensus sequence generation. The evolutionary history of SARS-CoV-2 was inferred using a Bayesian approach, implemented through the Markov chain Monte Carlo framework available in BEAST 1.10.4 ^31^, utilizing the BEAGLE library v3 ^32^ to increase computational performance. For poorly sampled locations, the Bayesian inference of unsampled genomes was associated with an epidemiologically informed sampling time and location, but not with observed sequence data. The phylogenetic tree was then plotted using the FigTree (v1.4.4) accompany with the BEAST package. The consensus genome sequences have not been deposited to any public database, pending institutional approval.

### Bioinformatics analysis

About 18,000 SARS-CoV-2 genome sequences were downloaded from NCBI on September 2^nd^, from which 500 incomplete sequences with less than two-thirds of the usual ~30,000 bp genome length were discarded. Each of the remaining 17,500 genome sequences was individually aligned to the reference SARS-CoV-2 genome (NC_045512.2) using the NEEDLE tool (Needleman-Wunsch global alignment of two sequences) from the EMBOSS package v6.6.0.0 ^33^. All nucleotide positions were based on the reference genome. Variants of SNPs were named using the SNP-sites software ^34^. Custom scripts in Perl, Python, and R were used for various calculations and graphics.

## ACKNOWLEDGEMENTS

I thank Dr. Matthew J. Binnicker, whose lab provided RNA extracts of COVID-19 samples. I also thank Drs. Gregory A. Worrell and Thomas P. Burghardt for their valuable comments. I especially thank my son Michael and my daughter Michelle who provided conceptual inputs to this work and proofreading on the initial draft of this manuscript.

No special funding was received for this work, although the author is supported by NIH BRAIN initiative R01 grant (NS112144 from NINDS and NIMH) and the Minnesota Partnership for Biotechnology and Medical Genomics (MNP #17.16). These funding sources had no role in the design of the study, in the collection/analysis/interpretation of the data, in the writing of the report, or in the decision to publish. The corresponding author had access to all of the data in the study and had final responsibility for the decision to submit this work.s

---

**Supplemental Table S1.** List of lineages containging inter-clade co-occurring SNP pairs.

| **Group 1** | **S** | **G** | **GH** | **GR** |
| --- | --- | --- | --- | --- |
| 11083 (V) | MT246475,<br>MT263395,<br>MT263416,<br>MT326050,<br>MT627229,<br>MT632538,<br>MT632798,<br>MT632803,<br>MT632947,<br>MT633004,<br>MT641506,<br>MT641526,<br>MT642142,<br>MT831341,<br>MT831378,<br>MT831833 |  | MT641485, MT642102, MT642184,<br>MT642352, MT683416, MT683417,<br>MT683418, MT821688, MT831376,<br>MT831383, MT831384, MT831405,<br>MT831570, MT831605, MT831630,<br>MT831645, MT831685, MT831688,<br>MT831700, MT831710, MT831725,<br>MT831745, MT831753, MT831783,<br>MT831786, MT831792, MT831796,<br>MT831810, MT831811, MT831818,<br>MT831826, MT831828, MT831834,<br>MT831835, MT831846, MT834032,<br>MT834206, MT834340, MT946870,<br>MT947554, MT947570, MT947574,<br>MT947590 | MT642111,<br>MT821586,<br>MT831134,<br>MT831170,<br>MT831323,<br>MT831380,<br>MT831406,<br>MT831599,<br>MT831663,<br>MT831789,<br>MT831807,<br>MT834167,<br>MT947555,<br>MT947560 |
| **Group 2** |  |  |  |  |
| 8782 (S) |  | <u>MT834063</u>,<br>MT834289 | MT642080, MT821570, MT834043 | <u>MT821583</u>,<br><u>MT834220</u> |
| 28144 (S) |  | MT833984,<br><u>MT834063</u>,<br>MT834337,<br>MT834360 | MT326065, MT642241 | <u>MT821583</u>,<br>MT834193,<br><u>MT834220</u> |
| **Group 3** |  |  |  |  |
| 25563 (GH) |  |  |  | MT831595 |

